# The Prevalence and Identification of Livestock Tick by Sex Ratio and Host in Tehran Province

**DOI:** 10.1101/2025.03.13.643006

**Authors:** Ebrahim Abbasi

## Abstract

Ticks are one of the most dominant forced ectoparasites of vertebrates, belonging to the arthropods, which transmit pathogens such as bacteria, viruses, and parasites to humans and animals in Iran and worldwide. Given that sex ratio factors can affect the epidemiology of vector-borne diseases, this study aimed to identify and determine the ticks’ sex ratio and host type (camels, sheep, cattle, dogs, chickens, and pigeons) in different areas of Tehran Province. This descriptive cross-sectional study took samples from different parts of the animal’s body in four seasons from 20 villages in 2019, in which 685 hard ticks and 121 soft ticks were caught from 1623 studied livestock and poultry. Regarding sex segregation among all caught ticks, 42.01% were male, and 57.99% were female. It is noteworthy that in both mountain and plain environments, *R. sanguineus sensu lato* species of hard ticks had the most elevated sex ratio. Most ticks were collected from sheep hosts, accounting for 60.04% of infestations, while the lowest infestation rate was found in cattle hosts at 0.62%.

## Introduction

Generally, ticks are divided into two large families, including Ixodidae (hard ticks) and Argasidae (soft ticks), with diverse species. They are the most critical obligate ectoparasites of animals, especially livestock and poultry (1). These blood-sucking ectoparasites, which are also known as pathogenic vectors, transmit bacteria, viruses, and protists to their hosts, including animals and humans. These pathogens cause various types of diseases, such as bacterial diseases (Q fever, Lyme disease, borreliosis, relapsing fever, and borreliosis), fungal diseases (dermatophilosis), protozoal diseases (babesiosis and theileriosis), and rickettsial diseases (ehrlichiosis, Brazilian spotted fever, anaplasmosis, and Rocky Mountain spotted fever) (2, 3).

Since sex ratio is a critical parameter determining the status and dynamics of animal populations, studies on sex ratio are vital for understanding the biology of populations and the biology of pathogens. Accordingly, arthropod vectors (e.g., ticks) and sex ratio could play different roles in pathogen transmission (4, 5).

Although ticks have been known since time immemorial, their role in causing livestock troubles was initiated in the mid-19th century. Due to the growth of the world’s population and nutritional needs, the number of livestock has increased rapidly throughout the industry. At the same time, concerns and issues related to ticks have emerged (6). In 1814, piroplasmosis was diagnosed in cattle in the United States. In 1821, it was discovered that the disease was transmitted to cattle by the ticks’ bite called *Rhipicephalus (Boophilus) annulatus* (7). In 1971, Mazloum conducted studies on the geographical distribution, seasonal activity, preferred tick hosts, and diseases transmitted to livestock and humans (8).

Pourmand et al. have also conducted a study to determine the frequency and species diversity of hard ticks and their sex ratio in equids in Sardasht suburb, West Azerbaijan Province, Iran. The results show that 85.48% of the ticks were male and 14.51% were female, with the highest frequency of *Hyalomma anatolicum* (67.74%), *Rhipicephalus bursa* (21.94%), H. marginatum (8.01%), and *Dermacentor marginatus* (2.29%), respectively (9).

Since tick bites can transmit diseases to livestock and poultry, it is important to identify the most common ticks based on the host and the sex ratio of the ticks, which can be a practical way to combat ticks and prevent the spread of diseases, ultimately minimizing economic losses due to livestock. Thus, this study aimed to determine the sex and identify ticks in different hosts, including camels, sheep, cattle, dogs, chickens, and pigeons in Tehran Province in 2019.

## Materials and Methods

### Geographical Area

This study was performed in 20 selected villages of Tehran Province with an area of approximately 185.956 square kilometers, located between 34 to 5.36 degrees north latitude and 50 to 53 degrees east longitude.

### Sampling Size and Method

The sample size was determined using the (10) procedure (d = 0.045, p = 0.3, (1-p) = 0.7). Accordingly, 685 hard ticks and 121 soft ticks were collected from 1623 livestock and poultry.

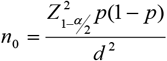

### Sample Collection and Identification

The feeding ticks on the animal body were separated from different parts of their bodies, such as earlobes, groin, tail base, and abdomen, and transferred to particular cans using a pence. Then, the specific characteristics of the animal (age, sex, livestock owner, and livestock code), village name, date of tick collection, collector name, and the number of caught ticks were written on the can and transferred to a unique flask to maintain the humidity and temperature required by the ticks.

The ticks were then transmitted to the laboratory for diagnosis and placed under a loop (stereomicroscope) to identify the sex and species using the valid diagnostic keys of the world and the region (11).

## Results

### Identification and Distribution of Livestock Ticks by Sex

This study gathered 685 hard ticks and 121 soft ticks from a total of 1623 domestic animals, including camels, sheep, cows, dogs, chickens, and pigeons infected with ticks. Regarding sex segregation among all captured ticks, the results showed that 42.01% of ticks were male, 57.99% female, and 15.01% were soft ticks.

In addition, among 685 hard-caught ticks (44.37%), 304 were male, and (55.62%) 381 were female. In both mountain and plain climates, *R. sanguineus sensu lato* has the highest sex ratio of hard ticks.

### Identifying and Determining the Distribution of Ticks by Host Type

Concerning tick-infested hosts, most ticks were collected from sheep (60.04%), and the lowest were collected from cattle (0.62%). Among the species of captured ticks in the family of hard ticks, the genus *Rhipicephalus (Boophilus)* was collected only from the cattle, the genus *Haemaphysalis* from the sheep and goats, and the genera *Ripisfalus* and *Hyalomma* were collected from all hosts except for pigeons, and chickens and in the corral wall. In the soft tick family, the genus *Ornithodoros* was collected only from the cage wall, and the genus *Argas* was collected from both pigeons and chickens. Unlike hard ticks, no soft ticks were caught from cattle, sheep, goats, camels, and dogs.

The frequency of tick species caught by host type is such that in cattle, small quantities of two species, *Hy. Marginatum* and *B. annulatus* were found. In sheep, *Rhipicephalus sanguineus sensu lato*, with 242 specimens, had the highest number, and *Rhipicephalus bursa*, with 5 specimens, had the lowest number. *R. sanguineus sensu lato* is found with the highest accumulation on the body of goats. In camels, the species *of Hy. Marginatum* had the highest frequency, and *R. sanguineus sensu lato* had the lowest frequency. In dogs, only *R. sanguineus sensu lato* with 19 numbers was found. In pigeons, only *A. reflexus* of the genus *Argas* has been collected. *A. persicus* was collected in significant abundance from the chickens’ bodies, and *O. lahorensis* was found only from the corral wall (Table 1 and Figure 1).

**Table 1.**
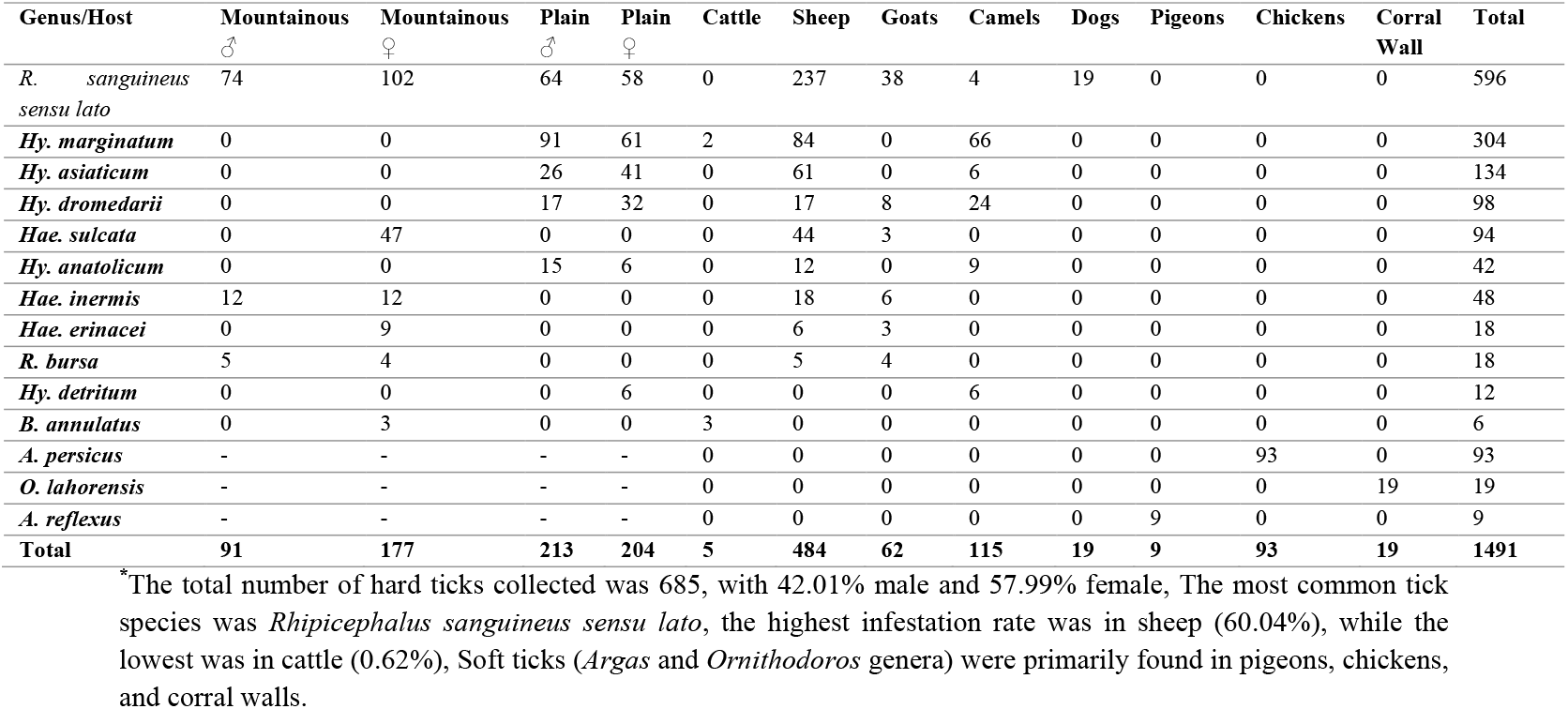
Comprehensive Identification and Distribution of Ticks in Tehran Province, 2019.

**Figure 1.**
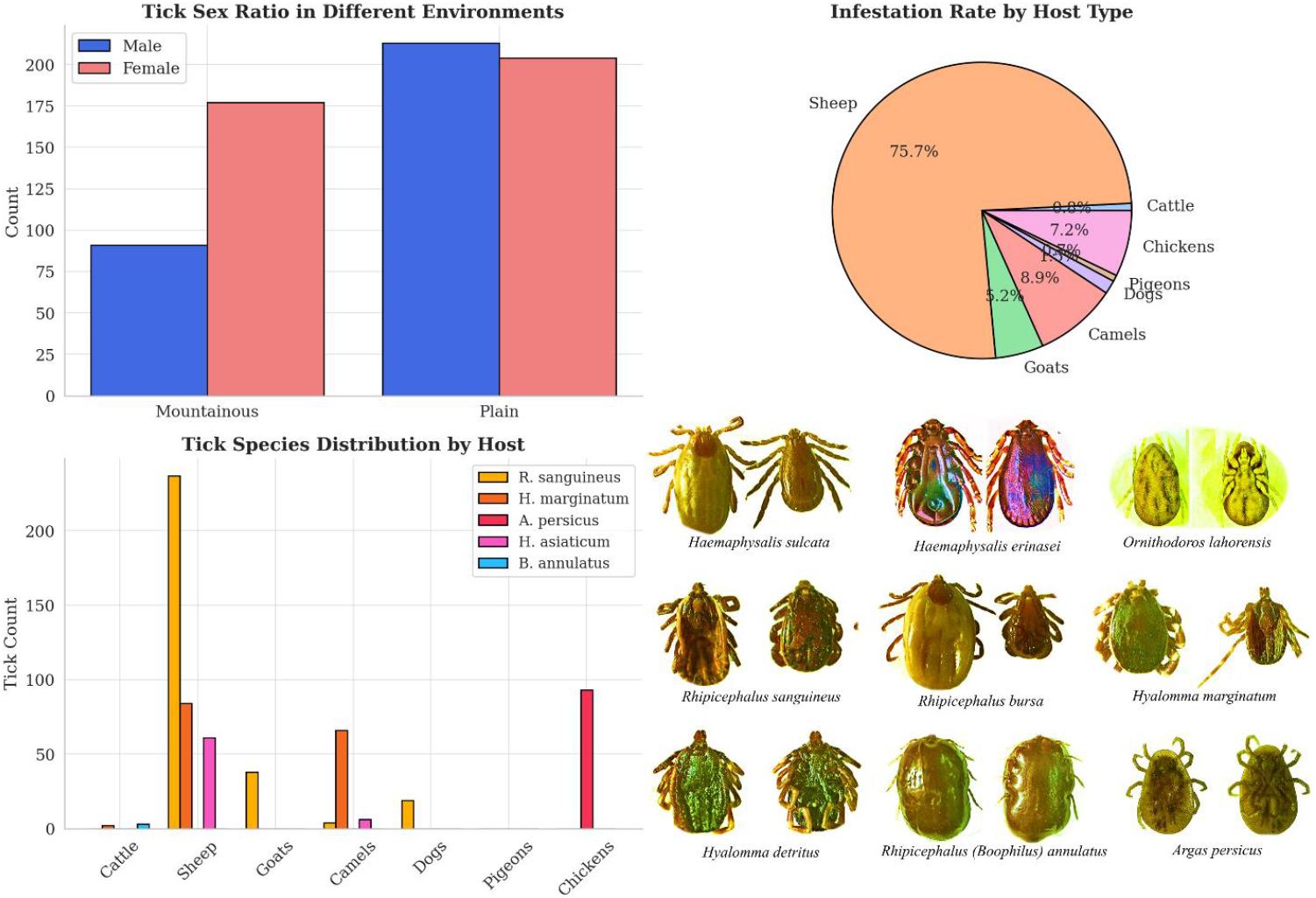
Bar Chart (Tick Sex Ratio in Different Environments): Sex-Based Distribution of Hard Ticks in Mountainous and Plain Environments, Pie Chart (Infestation Rate by Host Type): Proportion of Tick Infestation Among Different Livestock and Poultry, Grouped Bar Chart (Tick Species Distribution by Host): Host-Specific Distribution of Tick Genera in Tehran Province, collecting ticks from hosts and identifying them using loop (stereomicroscope).

## Discussion

The current study results indicate that sex ratios observed in ticks differ among species and even among host populations. Life-history aspects probably play a significant role in this regard. However, survival analyses under natural conditions are lacking in practically all tick genera, which are crucial to elucidating general patterns. Prospective molecular methods will provide new ways to determine the vast spectrum of possible performers affecting sex ratios. Previous studies demonstrated that sex ratios could depend on the season and area of collection. Also, in arthropods (e.g., ticks), sex ratios could be skewed towards females by reproductive parasites that appertain to this gender for their transmission (transovarial) (12, 13).

Our results showed that the highest rate of tick-infected livestock was related to sheep at 60.04%, and the lowest rate was related to cattle at 0.62% because the most studied livestock was sheep. The study in Ghaemshahr city demonstrated that the highest rate of infected livestock with ticks was observed in sheep at 28.3% and the lowest at 20% in cattle, which is compatible with our results and the lower infestation rates in cattle due to the smaller number of cattle included in the study compared to other livestock (14).

In addition, the dominant fauna species were found to be *Rhipicephalus Sanguineus*, which was consistent with the results of the majority of previous research (15, 16).

To conclude, the genus and species of dominant ticks in each region are diverse, and the geographical zone and climatic conditions of the area regulate the species and even the sex of active ticks in that province. Therefore, due to the high contamination of sheep, the authorities’ veterinary personnel and ranchers must be in control programs against external livestock parasites (mites) at least twice a year (with a maximum interval of 30 days), in addition to corral pesticide spraying, bathing the animals in the anti-tick bath.

## Declaration

### Ethics approval and consent to participate

Not applicable

### Data Availability Statement

All data generated or analyzed during this study are included in this published article.

### Competing interests

The authors declare no competing interests.

### Consent for publication

Not applicable

### Funding

This research received no specific grant from any funding agency in the public, commercial, or not-for-profit sectors.

### Authors’ contributions

E.A. has conducted all parts of the study, including design, execution, and writing the manuscript.

## Acknowledgments

The author would like to thank the Research Vice-Chancellor of Shiraz University of Medical Sciences.

